# Antibodies to native gluten arise from cross-reactive B cells with implications for epitope spreading in celiac disease

**DOI:** 10.1101/2022.01.29.478298

**Authors:** Chunyan Zhou, Thomas Østerbye, Shiva Dahal-Koirala, Øyvind Steinsbø, Jørgen Jahnsen, Knut E. A. Lundin, Søren Buus, Ludvig M. Sollid, Rasmus Iversen

## Abstract

Antibodies to deamidated gluten peptides are accurate diagnostic markers of celiac disease (CeD). However, antibody binding to all possible gluten epitopes has not previously been investigated. To map antibody reactivity in detail and to understand the connection between disease-relevant B-cell and T-cell epitopes, we took advantage of a high-density peptide array for assessment of serum antibody specificity in CeD across the wheat gluten proteome. We confirm the importance of peptide deamidation for antibody binding, and we show that the response is remarkably focused on the known epitope QPEQPFP (where E results from deamidation of Q). In addition, we describe a new epitope in native (non-deamidated) gluten, QQPEQII (where E is gene encoded), which was associated with both B-cell and T-cell reactivity. By generating monoclonal antibodies from peptide-binding gut plasma cells of CeD patients, we show that antibodies to this native gluten epitope are cross-reactive with the major deamidated epitope due to recognition of the shared PEQ motif. Hence, antibodies to native gluten appear to arise from cross-reactive B cells that are generated as a side effect of the immune response to deamidated gluten. Since cross-reactive B cells could present peptides to different gluten-specific T cells, we suspect that such B cells can play a role in epitope spreading by engaging T cells with multiple specificities.

## INTRODUCTION

Gluten is a collective term describing gliadin and glutenin protein families that are abundant in wheat flour. The term is also increasingly used to describe related proteins of rye (secalin), barley (hordein) and oats (avenin). Common for these proteins are high contents of the amino acids (aa) proline (P) and glutamine (Q) and a primary structure dominated by repetitive sequences^1^. In celiac disease (CeD), gluten proteins become the target of a harmful immune response that leads to tissue inflammation and characteristic morphological changes of the small intestinal mucosa^2^. Key pathogenic players are CD4+ T cells that recognize gluten peptides in the context of disease-associated HLA-DQ molecules. The majority of patients express HLA-DQ2.5, and the rest express either HLA-DQ2.2 or HLA-DQ8. Notably, conversion of certain gluten glutamine residues into glutamic acid (E) by deamidation stabilizes peptide binding to HLA and significantly increases T-cell reactivity^3-5^. In the body, deamidation of dietary gluten is mediated by the enzyme transglutaminase 2 (TG2), which specifically targets glutamine residues residing in QXP motifs (where X can be any amino acid)^6, 7^.

In CeD, there is autoantibody production against TG2 as well as antibodies against gluten. Both of these antibody responses are believed to rely on B-cell interaction with gluten-specific CD4+ T cells^8^. Similar to what was observed for T cells, gluten deamidation significantly increases the reactivity of CeD serum antibodies^9^. Through screening of individual gluten peptides, the sequence QPEQPFP (where E is introduced by deamidation of the TG2 target motif QQP) from γ- and ω-gliadin was identified as a common target of gluten-specific serum antibodies^9, 10^. The same sequence shows overlap with several T-cell epitopes, including the immunodominant DQ2.5-glia-ω1 and DQ2.5-glia-ω2^11^. It was also found to be recognized by most gluten-reactive plasma cells isolated from gut biopsies of CeD patients, indicating that it harbors a major B-cell epitope^12^. However, identification of this epitope was based on analysis of just a few gliadin proteins, raising the possibility that there could be additional disease-relevant gluten epitopes.

In order to map antibody binding to gluten peptides in detail and to understand the connection between B-cell and T-cell epitopes in CeD, we here utilized a high-density peptide array to analyze the specificity of CeD serum antibodies across the wheat gluten proteome. Peptides were synthesized in both their native and deamidated versions, allowing us to assess the effect of potential TG2-mediated deamidation on antibody binding. By using this approach, we confirm the importance of deamidation and the preference for the QPEQPFP epitope. In addition, we identify two new B-cell epitopes that dominate the response against native gluten peptides. Both of these contain gene-encoded PEQ motifs, and one of the sequences was also associated with T-cell reactivity. By generating monoclonal antibodies (mAbs) from gut plasma cells of CeD patients, we show that antibodies against this native gluten epitope are cross-reactive with deamidated peptides through recognition of the shared PEQ motif. In CeD, B cells with such cross-reactive B-cell receptors (BCRs) may facilitate epitope-spreading by serving as antigen-presenting cells (APCs) for T cells reactive with different gluten peptides.

## RESULTS

### Identification of antibody epitopes in gluten

In order to scan through the gluten proteome for antibody reactivity, we generated overlapping peptides covering 904 individual gluten proteins identified in wheat. The peptides were synthesized on a high-density peptide array as 15mers with 14 aa overlap, resulting in display of 36,231 unique peptide sequences. The peptides were incubated with pooled serum samples of ten CeD patients or two non-celiac control donors (Table S1), followed by detection of bound IgA or IgG antibodies. While no reactivity was observed in the control sera, a number of peptides were recognized by IgA and IgG antibodies in CeD patients (Table S2). Among the IgA antibodies, there was a strong preference for binding of peptides containing the TG2 target motif QXP (Fig. 1A). This selectivity could also be observed for the peptides with the highest IgG reactivity, indicating that both the IgA and IgG anti-gluten response in CeD are biased toward peptides that can be deamidated by TG2.

**Figure 1.**
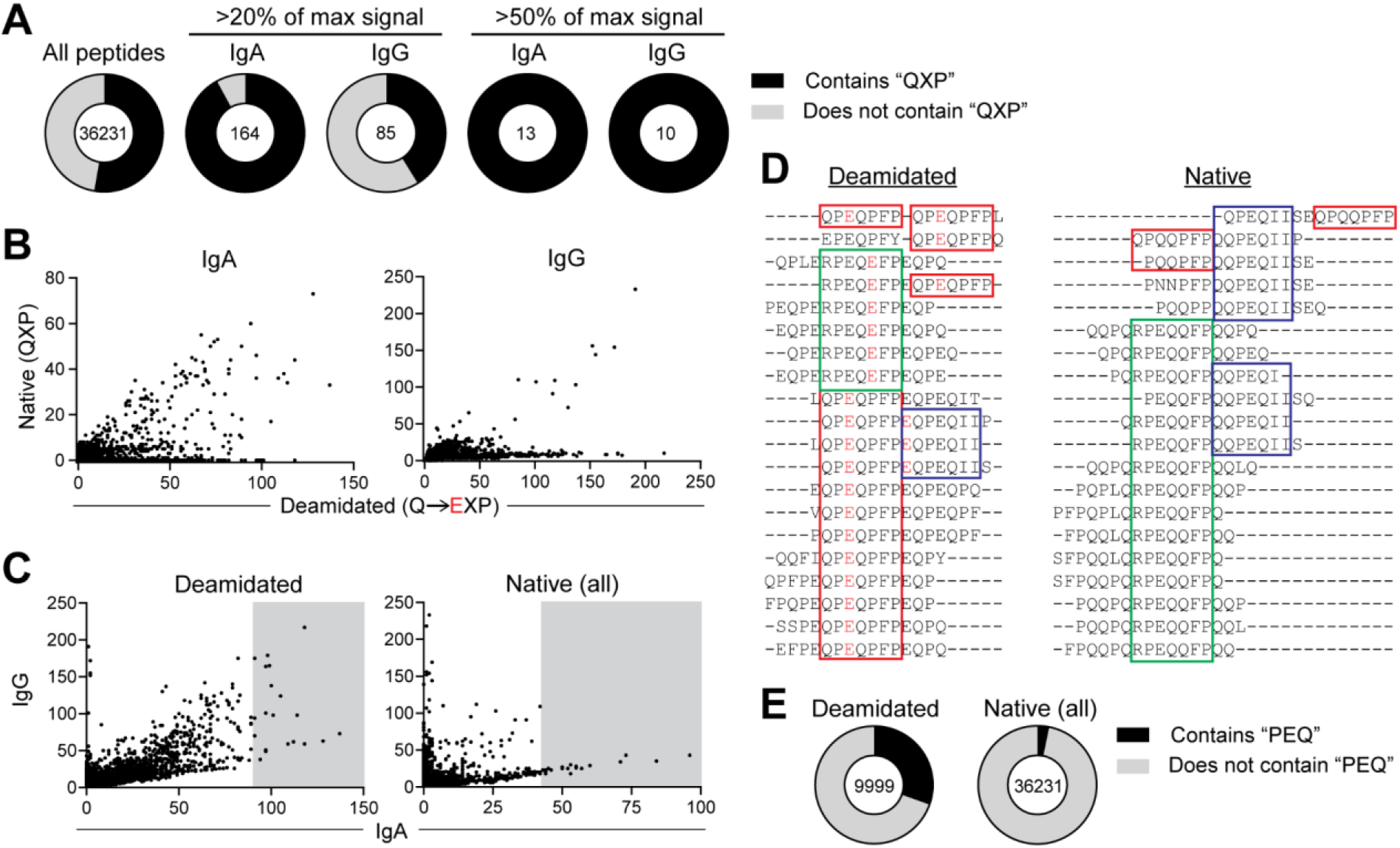
Serum antibody reactivity to gluten peptides. (A) Occurrence of the “QXP” motif in unique 15mer gluten peptides synthesized on a high-density peptide array. The number of peptides associated with high IgA and IgG reactivity in pooled sera of CeD patients is indicated. (B) Antibody reactivity to individual peptides that were synthesized in both their native (QXP) and deamidated (EXP) form. (C) Relationship between IgA and IgG reactivity to individual deamidated and native gluten peptides. The top-20 IgA-reactive peptides are indicated with a grey box. (D) Sequence alignment of the top-20 IgA-reactive deamidated and native peptides generated with the Clustal Omega multiple sequence alignment tool13. Three 7 aa epitopes present in both deamidated and native peptides are indicated with individual colored boxes. Within these epitopes, glutamic acid residues resulting from deamidation are indicated in red. (E) Occurrence of the “PEQ” motif in deamidated and native gluten peptides that were represented on the array.

To assess the role of deamidation, peptides containing the QXP motif were also synthesized in their deamidated (EXP) form (Table S3). All peptides that were recognized by antibodies in their native state were also recognized in their deamidated version (Fig. 1B). For most of these peptides, deamidation increased antibody reactivity. Moreover, many peptides that were not recognized in their native form showed reactivity upon deamidation. These results thus confirm that there is strong selection for deamidated epitopes among anti-gluten antibodies in CeD.

When comparing the IgA and IgG reactivity to individual peptides, we observed that some peptides were uniquely targeted by IgG (Fig. 1C and Fig. S1). In general, however, the peptides with high IgA reactivity also showed an IgG signal, and we decided to focus on these in our further analysis. By aligning the sequences of the top-20 IgA-reactive native and deamidated peptides, we identified three 7 aa epitope motifs that collectively could explain all of the antibody reactivity (Fig. 1D). Among the deamidated peptides, 15 of the 20 sequences contained the previously identified epitope QPEQPFP, confirming its dominant role in the anti-gluten response. The non-deamidated version of this sequence was also observed in a few of the targeted native peptides. However, two other sequences, RPEQQFP and QQPEQII, dominated among native gluten peptides with high IgA reactivity. Curiously, all three identified epitopes share the short motif PEQ. This motif is found in 30% of all deamidated peptides but only in 3% of the native gluten peptides (Fig. 1E). Thus, our analysis of CeD serum antibody reactivity demonstrates a remarkable selection for gluten peptides carrying the PEQ motif.

### Native gluten epitope is recognized by T cells

The two native gluten epitopes, we describe here, are found in an ω-secalin protein, which was identified in wheat cultivars carrying a rye translocation^14^. Further, one of the sequences shows high similarity to a previously identified T-cell epitope of rye ω-secalin^15^ (DQ2.5-sec-3, Fig. 2A). To understand if the two newly identified antibody epitopes are also recognized by T cells in CeD, we generated overlapping peptides carrying one or both of the sequences and tested reactivity among a panel of gluten-reactive T-cell lines obtained from CeD patient gut biopsies. One out of 16 lines showed reactivity, and only when the DQ2.5-sec-3-related sequence was present (Fig. 2B). These results demonstrate that one of the identified antibody epitopes in native gluten is also associated with T-cell reactivity, although T cells with this specificity are likely rare in CeD.

**Figure 2.**
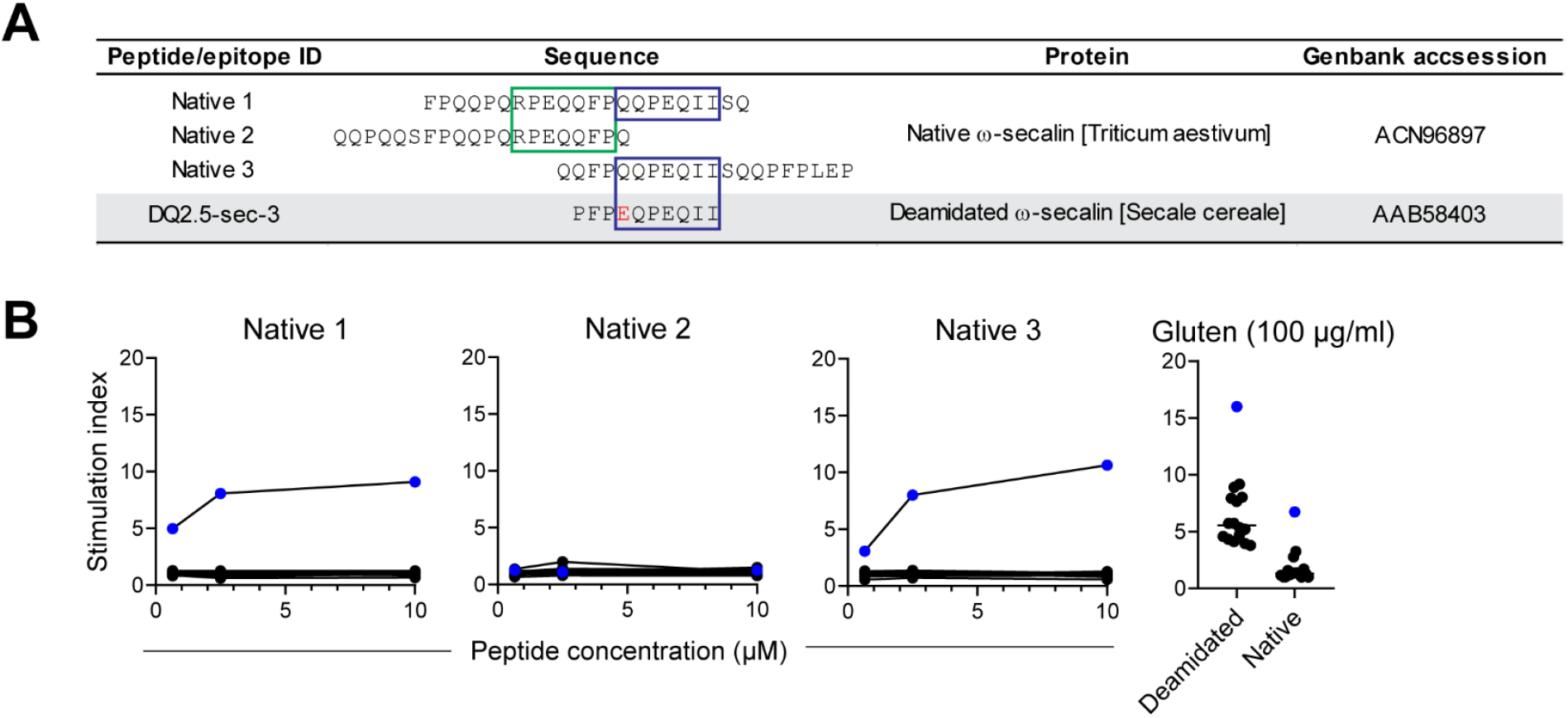
T-cell reactivity to native gluten epitopes. (A) Overview of overlapping peptides used to represent the identified antibody epitopes in native wheat (*Triticum aestivum*) gluten and comparison with the previously reported T-cell epitope DQ2.5-sec-3 in rye (*Secale cereale*). A glutamic acid residue introduced by deamidation in the DQ2.5-sec-3 epitope is indicated in red. (B) Reactivity of 16 individual T-cell lines obtained from gut biopsies of CeD patients assessed by thymidine incorporation. The T cells were stimulated with the indicated peptides or chymotrypsin-digested gluten that was either added directly or pre-treated with TG2 to induce deamidation. The stimulation index was calculated as the signal obtained with antigen divided by the signal obtained with PBS only. Horizontal lines indicate medians. One T-cell line that showed reactivity to peptides containing the QQPEQII sequence is indicated with blue symbols.

### Serum antibody cross-reactivity to deamidated and native epitopes

Since we could only confirm T-cell reactivity against one of the native antibody epitopes, we decided to focus on this sequence in our further analysis of the anti-gluten immune response. To compare antibody reactivity against the major deamidated and native epitopes (referred to in the following as “anti-deamidated” and “anti-native” antibodies), we assessed binding of synthetic peptides to serum IgA of individual CeD patients or non-celiac control donors (Fig. 3A). For both epitopes, binding was specific to CeD, and patients expressing HLA-DQ2.5 generally displayed higher reactivity than HLA-DQ8 patients, in agreement with an overlap between the identified antibody epitopes and HLA-DQ2.5-resticted T-cell epitopes. Antibody reactivity was highest to the deamidated epitope, and many patients did not have detectable levels of anti-native antibodies. Notably, all patients with antibodies to the native epitope also had reactivity to the deamidated epitope, suggesting that formation of anti-native antibodies might depend on the anti-deamidated response (Fig. 3B).

**Figure 3.**
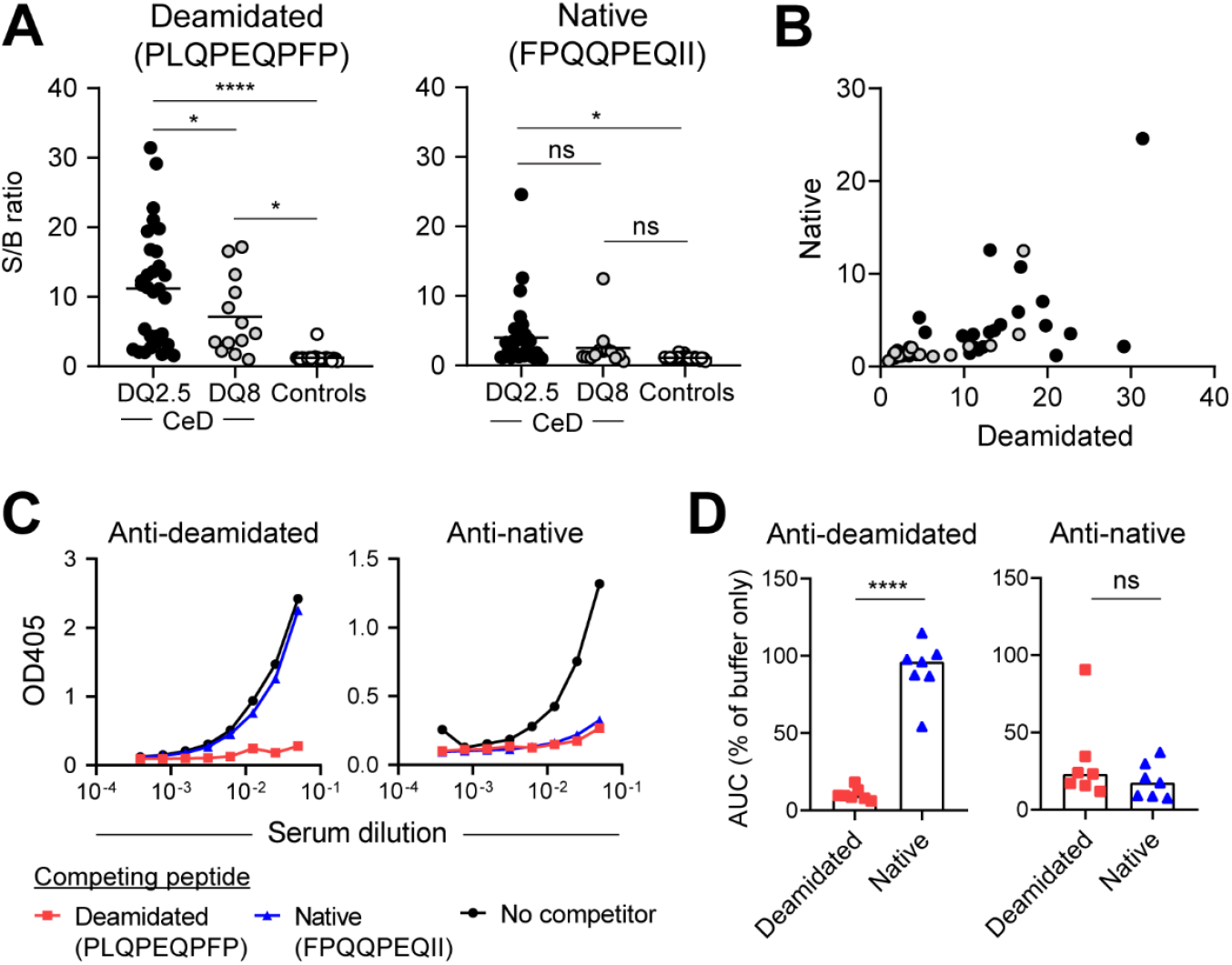
IgA reactivity to deamidated and native gluten epitopes in individual serum samples. (A) IgA reactivity to peptides representing the major deamidated and native epitopes in individual serum samples (1:20 dilution) of CeD patients expressing HLA-DQ2.5 (n = 30) or HLA-DQ8 (n = 13) or in non-celiac controls (n = 24) assessed by ELISA. The signal-to-background (S/B) ratio was calculated based on the signal obtained without coating of peptide. Horizontal lines indicate means, and differences between groups were analyzed by one-way ANOVA with Holm-Sidak multiple comparisons correction. \**P* < 0.05; **** *P* < 0.001. (B) Correlation between serum IgA reactivity against the deamidated and native epitopes in individual CeD patients. Colors represent HLA-DQ variants as shown in (A). (C and D) Effect of pre-incubation with immobilized competing peptide prior to assessment of IgA reactivity to the deamidated or native epitopes. Serum titration curves are shown for a representative CeD patient in (C). Titration curves obtained from seven CeD patients were used to calculate area-under-the-curve (AUC) values plotted in (D). Bar heights indicate means, and statistical difference was evaluated by a paired *t* test.

To test if the observed anti-gluten binding pattern could be explained by antibody cross-reactivity, we pre-incubated serum samples of CeD patients with immobilized peptides harboring the deamidated or native epitopes and assessed remaining IgA reactivity in the supernatant (Fig. 3C-D). The native peptide did not affect binding to the deamidated epitope, but pre-incubation with the deamidated peptide efficiently abrogated binding to the native epitope. These results indicate that most anti-deamidated antibodies are specific, whereas antibodies binding to the native epitope are cross-reactive with the deamidated epitope.

### Cross-reactivity of gut plasma cells binding to native epitope

To see if we could identify plasma cell populations corresponding to the anti-gluten antibodies in serum, we stained duodenal single-cell suspensions of CeD patients with the native and deamidated peptides and assessed binding by flow cytometry (Fig. 4A). In agreement with previous results, we could detect a population of surface IgA+ plasma cells that recognized the deamidated peptide^12, 16, 17^. In addition, we could detect a smaller population of cells that bound the native peptide. These cells made up less than 0.3% of IgA+ gut plasma cells, mirroring the relatively low serum IgA reactivity observed in most patients. To further characterize the mucosal antibody response, we sorted single native-reactive gut plasma cells (Table S1) and generated recombinant mAbs by expression cloning. Of 19 tested mAbs, 17 were confirmed reactive with the native peptide. In addition, most of the mAbs showed almost equal reactivity to the deamidated peptide (Fig. 4B-C and Table S4). These results thus corroborate our serum analysis and confirms that antibodies binding to the identified native gluten epitope are cross-reactive with the major deamidated epitope.

**Figure 4.**
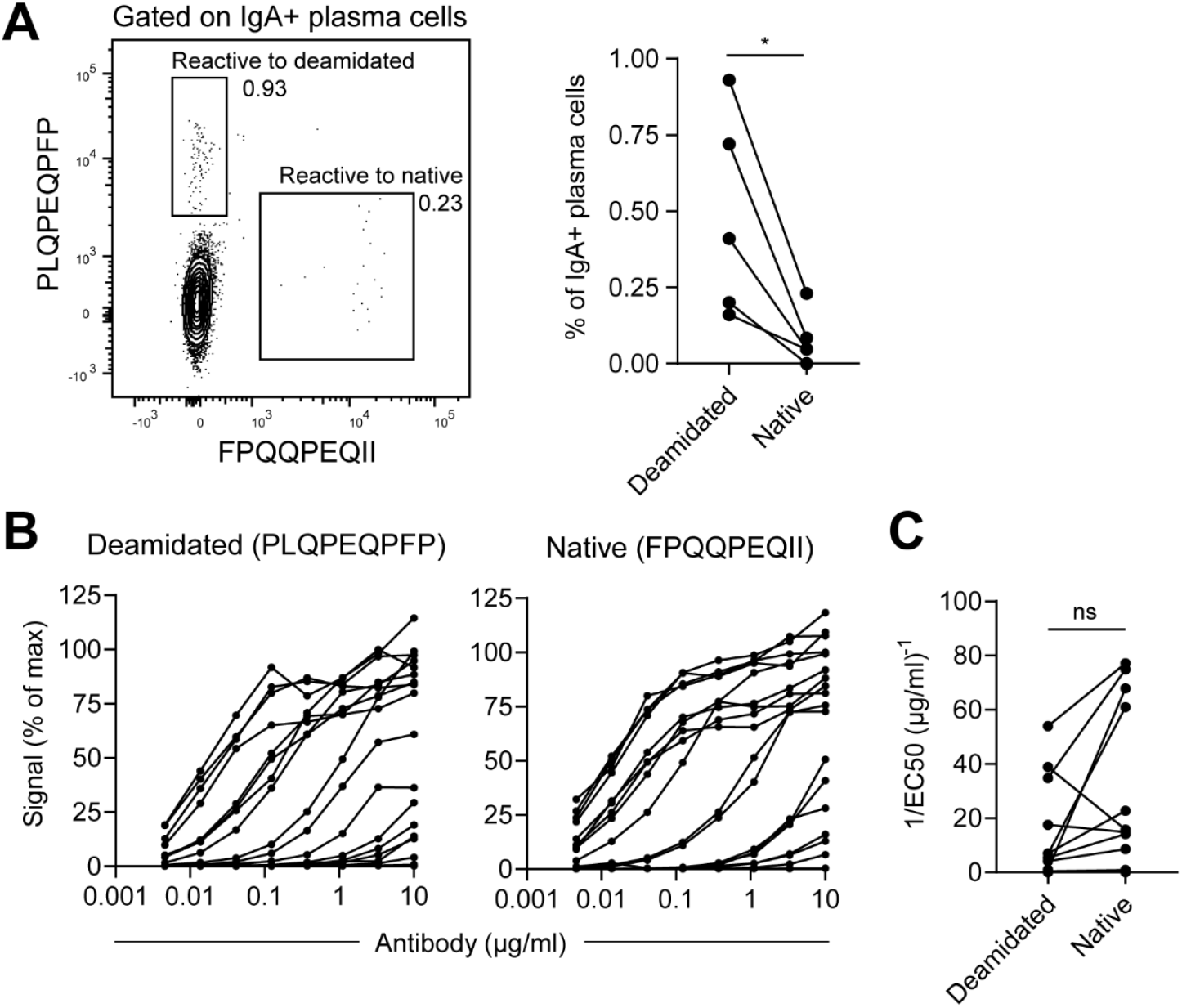
Generation of mAbs from native-reactive gut plasma cells. (A) Representative flow cytometry plot and summary data of five CeD patients showing staining of IgA+ gut plasma cells with peptides representing the deamidated and native epitopes. (B and C) Reactivity of 19 mAbs generated from gut plasma cells to the deamidated and native epitopes in ELISA. Based on titration binding curves (B), EC50 values were calculated and used to compare the affinity to the deamidated and native epitopes for 17 confirmed native-reactive mAbs (C). Differences between groups were analyzed by a paired *t* test. \**P* < 0.05.

### Antibodies binding to native gluten epitope have characteristic sequences

Many of the native-reactive mAbs used the heavy chain V-gene segment *IGHV3-15* in combination with the kappa light chain segment *IGKV4-1* (Fig. 5A). Plasma cells with these V-gene segments also showed signs of clonal expansion in CeD patients (Table S4), indicating strong selection for a particular type of antibodies in the response against native gluten. The same heavy and light chain combination was also found to dominate among gut plasma cells reactive with the major deamidated epitope^12, 16, 17^. However, the native-reactive mAbs tended to have longer CDR-H3 loops (Fig. 5B), which were characterized by a conserved amino acid sequence derived from the D-gene segment *IGHD3-10* (Fig. 5C and Table S4). Thus, antibodies reactive with the native epitope have particular features that distinguish them from antibodies to the deamidated epitope. In line with this notion, and in agreement with our serum IgA analysis, anti-deamidated mAbs using the stereotypical V-gene segments did not show strong cross-reactivity to the native epitope (Fig. 5D).

**Figure 5.**
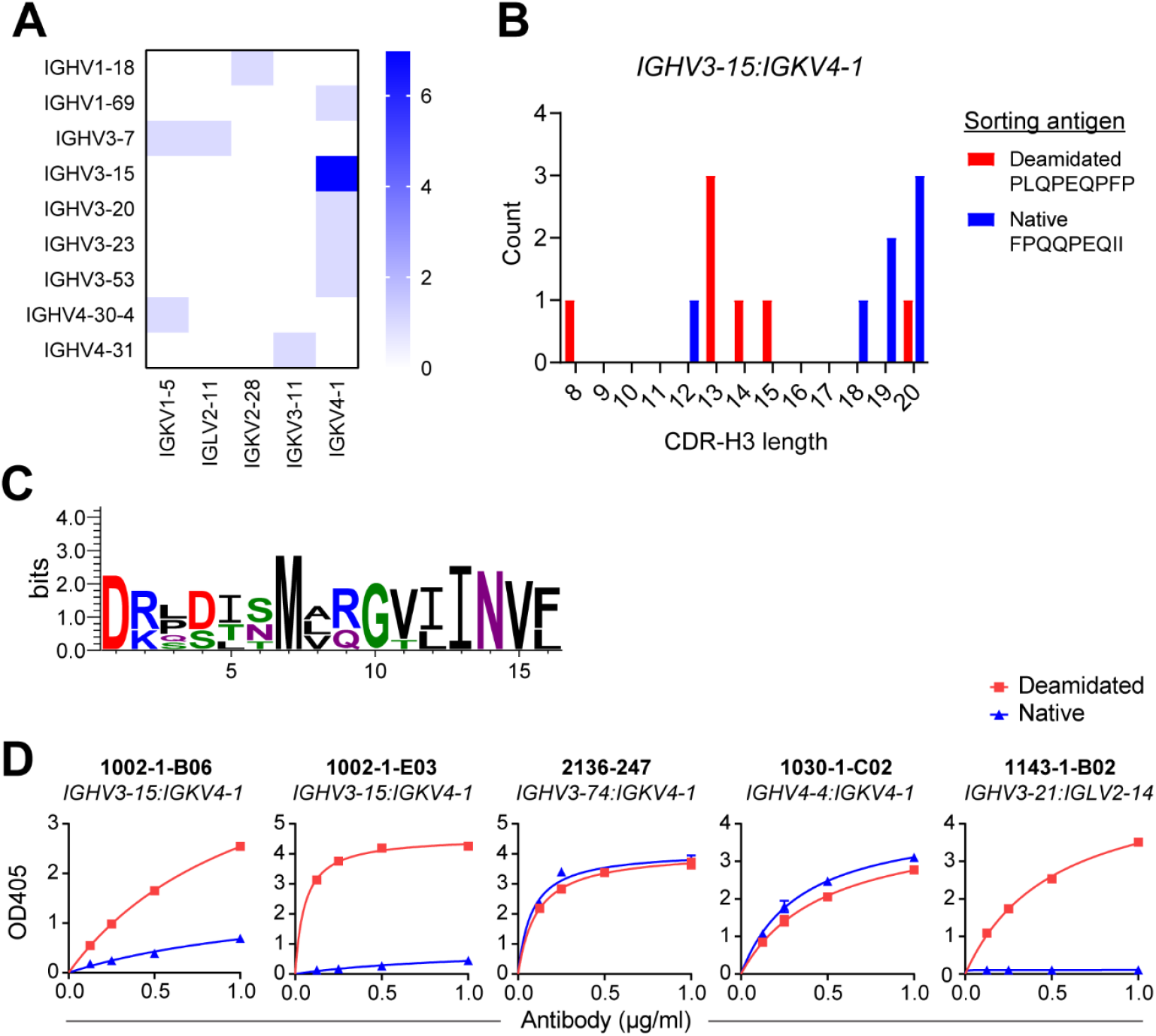
Characteristics of native-reactive mAbs. (A) Heatmap showing pairing of heavy and light chain V-gene segments among 16 clonotypes reactive with the native gluten epitope. (B) CDR-H3 aa length of clonotypes using *IGHV3-15* in combination with *IGKV4-1*. Sequences were obtained from plasma cells reactive with the native or deamidated^12^ gluten epitope. (C) Sequence logo showing the core CDR-H3 sequence of native-reactive *IGHV3-15:IGKV4-1* plasma cells with 19 or 20 aa long CDR-H3 loops. The logo was compiled from six sequences using WebLogo v3.7.4^18^. (D) ELISA titration curves showing binding of mAbs generated from deamidated-reactive plasma cells12 to either the deamidated or native gluten epitope. Two mAbs using *IGKV4-1* in combination with non-canonical *IGHV* segments show cross-reactivity, whereas two stereotypical *IGHV3-15:IGKV4-1* mAbs only bind weakly to the native epitope.

### Cross-reactivity of anti-gluten BCRs

Gluten- and TG2-reactive B cells are believed to play a key role as APCs for pathogenic CD4+ T cells in CeD^8, 19, 20^. Since BCR specificity determines antigen uptake and presentation, we generated transduced A20 B-cell lines expressing selected stereotypic mAbs in the form of membrane-bound BCRs in order to address the ability of gluten-reactive B cells to bind different peptides. B cells representing the deamidated and native epitope reactivities were both able to bind several gluten-derived peptides in flow cytometry (Fig. 6A). In both cases, however, binding was completely dependent on the PEQ motif. Thus, the B cells did not recognize a major T-cell epitope from α-gliadin (DQ2.5-glia-α2, containing PEL), nor did they bind to the non-deamidated version of γ-gliadin (containing PQQ). In agreement with our analysis of soluble antibodies, the anti-native BCR showed a higher degree of cross-reactivity than the anti-deamidated BCR (Fig. 6B). Consequently, when incubated with a mixture of the deamidated and native peptides, the native-reactive B cells bound both peptides simultaneously, whereas the deamidated-reactive B cells only showed binding to the deamidated peptide (Fig. 6C).

**Figure 6.**
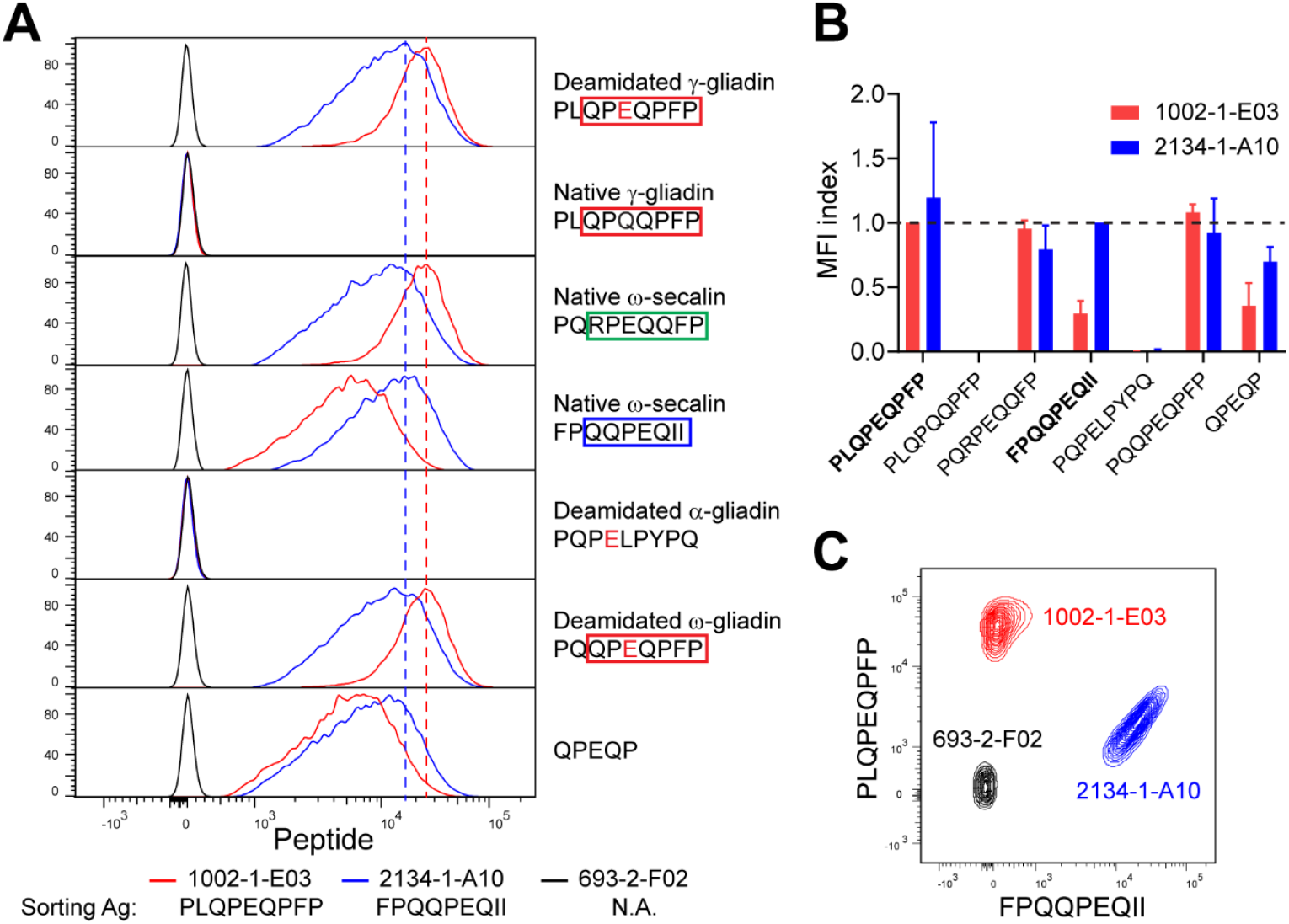
Binding of BCRs to gluten peptides. (A) Flow cytometry histograms showing binding of BCR-transduced A20 B cells to different gluten peptides. BCRs were generated from plasma cells reactive with the deamidated (1002-1-E03^16^) or native (2134-1-A10) gluten epitope or from a negative control with unknown specificity (693-2-F02^21^, N.A. not applicable). For each of the gluten-reactive BCRs, the dashed line indicates the peak of the signal obtained with the peptide antigen (Ag) that was used for plasma cell sorting. (B) Quantification of staining obtained with individual peptides. The deamidated and native peptides that were used for plasma cell sorting are indicating in bold, and the median fluorescence intensity (MFI) index was calculated as the ratio between MFI values obtained with each peptide and the peptide that was used for sorting. Error bars indicate SD based on 2 or 3 independent experiments. (C) Flow cytometry plot showing double staining of the A20 B cells with peptides representing the major deamidated and native epitopes.

### Cross-reactive B cells can present peptides to T cells with different specificities

To understand the implications of BCR cross-reactivity for interactions between T cells and B cells in CeD, we generated TCR-transduced BW58 hybridoma T-cell lines specific to either DQ2.5-glia-ω2 or DQ2.5-sec-3, corresponding to the major deamidated and native B-cell epitopes (Fig. 7A). When incubated with TG2-treated chymotrypsin-digested gluten, A20 B cells expressing native-reactive BCR in combination with HLA-DQ2.5 were able to stimulate both T cells, whereas B cells with a deamidated-reactive BCR only activated the DQ2.5-glia-ω2-specific T cells (Fig. 7B). These results demonstrate that gluten-reactive B cells with the ability to bind multiple peptides can engage T cells with different specificities.

**Figure 7.**
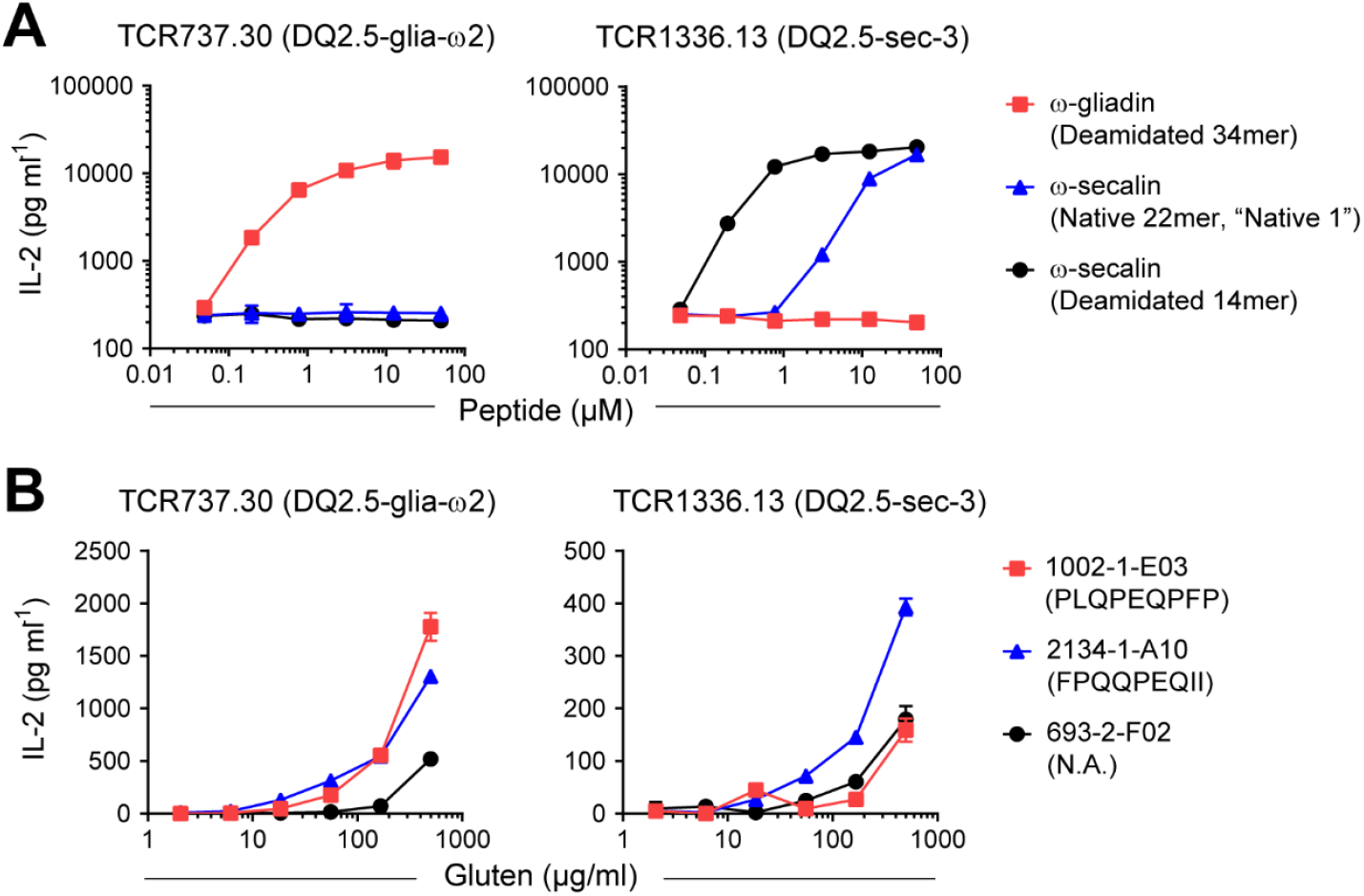
Presentation of peptides to gluten-reactive T cells. (A) Activation of TCR-transduced BW58 T cells measured as release of IL-2 after incubation with HLA-DQ2.5-expressing A20 B cells and synthetic gluten peptides. DQ2.5-glia-ω2-specific and DQ2.5-sec-3-specific T cells show the expected reactivity to deamidated ω-gliadin and ω-secalin, respectively. In addition, DQ2.5-sec-3-specific T cells show some reactivity to a peptide harboring the native B-cell epitope (Native 1, Fig. 2). (B) T-cell activation after incubation with a TG2-treated gluten digest and A20 B cells expressing HLA-DQ2.5 in combination with gluten-reactive (1002-1-E03 and 2134-1-A10) or non-gluten-reactive (693-2-F02) BCRs. Peptides used for plasma cell sorting and generation of recombinant BCRs are indicated (N.A. not applicable). The figure shows data from one of two experiments, and error bars indicate SD based on culture duplicates.

## DISCUSSION

So far, 27 HLA-DQ2.5-resticted gluten T-cell epitopes have been described in CeD^22^. All of these contain TG2-targeted deamidation sites, and deamidation significantly improves the reactivity of patient-derived T-cell clones. Four immunodominant gliadin epitopes (DQ2.5-glia-α1, DQ2.5-glia-α2, DQ2.5-glia-ω1 and DQ2.5-glia-ω2) are recognized across patients and are collectively responsible for around half of the anti-gluten T-cell reactivity in the gut^11, 23^. The gluten-reactive T cells are believed to interact with both TG2-specific and gluten-specific B cells, leading to mutual T-cell and B-cell activation and production of both TG2-specific and gluten-specific antibodies^8^. In order to understand the connection between gluten T-cell and B-cell epitopes, we here took advantage of a high-density peptide array to scan through the wheat gluten proteome for antibody reactivity. As seen for T-cell reactivity, peptide deamidation led to improved antibody binding. The antibody response to deamidated peptides was remarkably focused toward the sequence QPEQPFP, which was previously described as an important B-cell epitope in CeD^9, 10, 12, 24^. In addition, we here describe two new B-cell epitopes in native (non-deamidated) gluten, RPEQQFP and QQPEQII, of which the latter was also associated with T-cell reactivity.

By analyzing both polyclonal serum IgA and mAbs generated from gluten-reactive gut plasma cells, we show that binding to the QQPEQII epitope is mediated by cross-reactive antibodies that also bind to the major deamidated epitope through recognition of the shared core motif PEQ. B cells displaying this type of cross-reactivity were able to activate T cells with different specificities through BCR-mediated uptake and presentation of diverse peptides in a complex gluten digest. Such B cells may thus play a role in T-cell epitope spreading through presentation of multiple gluten peptides to T cells. In particular, we show that cross-reactive B cells can interact with both DQ2.5-glia-ω2-specific and DQ2.5-sec-3-specific T cells. We therefore speculate that T cell-B cell interactions involving immunodominant DQ2.5-glia-ω2-specific T cells may lead to accumulation of cross-reactive B cells that can go on to activate rare DQ2.5-sec-3-specific T cells.

Preferential targeting of the deamidated sequence QPEQPFP indicates that the anti-gluten antibody response is biased toward γ- and ω-gliadin proteins. The B-cell response is thus linked to the immunodominant DQ2.5-glia-ω1 and DQ2.5-glia-ω2 T-cell epitopes. However, the two other immunodominant T-cell epitopes, DQ2.5-glia-α1 and DQ2.5-glia-α2, were not major targets in the B-cell response, as we did not observe notable antibody reactivity to α-gliadin peptides. A possible explanation for this apparent discrepancy is that T cells specific to DQ2.5-glia-α1 and DQ2.5-glia-α2 primarily interact with TG2-specific rather than gluten-specific B cells. Thus, the anti-gluten antibody response might be shaped by competition between gluten-specific and TG2-specific B cells for T-cell help. TG2-specific B cells can interact with gluten-specific T cells through BCR-mediated uptake of TG2-gluten enzyme-substrate complexes^25^. TG2-specific B cells are therefore expected to primarily present peptides that form the most stable complexes with TG2. In support of a scenario where TG2-specific B cells are dedicated to presentation of α-gliadin peptides, the DQ2.5-glia-α1 and DQ2.5-glia-α2 epitope sequences were previously identified as preferred TG2 substrates in a complex wheat gluten digest^26^.

In summary, we have shown that the anti-gluten antibody response in CeD is highly focused toward the deamidated epitope QPEQPFP in γ- and ω-gliadin proteins. In some patients, recognition of the PEQ motif facilitates formation of cross-reactive antibodies that can bind PEQ-containing sequences in both native and deamidated gluten proteins. B cells showing this type of cross-reactivity can efficiently present peptides to T cells with different specificities, and they may thereby mediate spreading of the T-cell response to include rare epitopes such as DQ2.5-sec-3.

## MATERIALS AND METHODS

### Patients

Biological samples were obtained from CeD patients and non-celiac controls who had given their informed consent. The diagnosis of CeD was established according to the guidelines of the European Society for the Study of Coeliac Disease^27^. Ethical approval for the experimental protocols was obtained from the Regional Ethics Committee of South-Eastern Norway (REK ID 6544).

### High-density peptide array

A database of the wheat gluten proteome was compiled by extracting *Triticum aestivum* entries from the UniProt Knowledgebase^28^. To represent all possible gluten epitopes, 15mer peptides having 14 aa overlap with neighboring sequences were synthesized on a high-density peptide array as previously described^29^. Peptides harboring QXP motifs were also synthesized in their deamidated (EXP) version. To assess binding of serum antibodies, the chip was incubated with pooled sera of CeD patients or non-CeD controls diluted 1:200 in 50 mM Tris-acetate pH 8.0, 130 mM NaCl, 0.1% (v/v) Tween-20, 1 mg/ml BSA followed by detection of bound IgA and IgG with goat anti-human IgA-Cy5 and goat anti-human IgG-Cy3, respectively.

### T-cell proliferation assay

Polyclonal T-cell lines obtained from gut biopsies of CeD patients were used for setting up a ^3^H-thymidine incorporation-based T-cell proliferation assay as described previously^30^. In brief, 75,000 APCs (EBV-immortalized B-LCL from an HLA-DR3, DQ2.5 homozygous donor) were irradiated (75 Gy) and incubated at 37°C with different concentrations (10 μM, 2.5 μM and 0.0625 μM) of synthetic peptides (GenScript) or chymotrypsin-digested gluten (100 μg/ml, with or without TG2 treatment to introduce deamidation) or PBS. After 24 h, 50,000 T cells from the individual T-cell lines were added. The cultures were pulsed with 1 μCi ^3^H-thymidine 48 h later. T-cell proliferation was measured by ^3^H-thymidine incorporation with readout of CPM (counts per minute) 16-20 h after ^3^H-thymidine pulsing. All culture conditions were performed in triplicates. The T-cell lines were considered reactive if the stimulation index (CPM antigen/CPM PBS) was higher than 3.

### Flow cytometry and single-cell sorting

BCR binding to gluten peptides was assessed using tetramerized synthetic peptides (obtained from GenScript or GL Biochem) containing an N-terminal biotin moiety and a GSGSGS linker. Peptide tetramers were generated by incubating PE-conjugated streptavidin (Thermo) with individual peptides at a molar ratio of 1:1.5. When staining with two peptides simultaneously, FPQQPEQII was attached to streptavidin-PE, and PLQPEQPFP was attached to streptavidin-APC (Agilent). After incubation, unoccupied binding sites were blocked with free biotin, before the tetramers were added to the cells. The level of staining was analyzed by flow cytometry using an LSRFortessa instrument (BD). For sorting of peptide-reactive gut plasma cells, duodenal biopsies of CeD patients were processed into single-cell suspensions as previously described^31^ followed by staining with tetramerized peptides in combination with the following antibodies: Anti-human CD38-FITC (clone HB7, eBiosciences), anti-human IgA-APC/Vio770 (clone IS11-8E10, Miltenyi), anti-human CD3-Brilliant Violet 510 (clone OKT3, BioLegend), anti-human CD14-Brilliant Violet 510 (clone M5E2, BioLegend). Single IgA+ plasma cells (identified as large CD3^−^CD14^−^CD38^hi^ lymphocytes) binding to FPQQPEQII were then sorted into 5 μl of 1% (v/v) Nonidet P40 Substitute, 20 mM Tris-HCl, pH 8.0, containing 5 U murine RNase Inhibitor (NEB) using a FACSAriaIII instrument (BD). After sorting, cell lysates were flash-frozen on dry ice and stored at −80°C.

### Generation of recombinant mAbs

Heavy and light chain variable regions were amplified from single-cell lysates by a nested RT-PCR approach using previously reported V gene-specific forward primers^32^ in combination with IgA-specific reverse primers (3’CαCH1, TGGGAAGTTTCTGGCGGTCACG; 3’CαCH1-2, GTCCGCTTTCGCTCCAGGTCACACT). The PCR products were sequenced and cloned into vectors for expression of full-length human IgG1 in HEK293-F cells as previously described^16^. Sequence analysis was done with the IMGT/V-QUEST tool^33^.

### ELISA

To assess binding of serum antibodies and recombinant mAbs to the deamidated and native epitopes, synthetic biotinylated peptides were attached to streptavidin-coated ELISA plates (Thermo) using a peptide concentration of 50 nM in TBS with 0.1% (v/v) Tween-20 (TBST). After washing, the coated peptides were incubated with sera or recombinant mAbs diluted in 3% (w/v) BSA/TBST. Bound antibodies were subsequently detected with AP-conjugated goat anti-human IgA (Sigma) or goat anti-human IgG (Southern Biotech).

### Competition ELISA

To assess cross-reactivity of serum antibodies to the deamidated and native epitopes, synthetic biotinylated peptides were first attached to streptavidin-coated M-280 Dynabeads (Thermo) using 40 pmol of peptide and 20 μl beads in TBST. The beads were then washed using a magnetic separator and incubated with individual serum dilutions in TBST. The supernatant was collected and used for assessment of IgA binding to the native and deamidated epitopes in ELISA as described above. Signals obtained with and without incubation with peptide-coated beads were compared in order to determine the ability of each competitor peptide to adsorb antibody reactivity.

### Generation of transduced B-cell and T-cell lines

Antigen-specific B-cell and T-cell lines were generated by retroviral transduction of A20 lymphoma B cells expressing HLA-DQ2.5 and BW58α^−^β^−^hybridoma T cells expressing human CD4, respectively^21, 34^. To generate FPQQPEQII-reactive B cells, we obtained synthetic DNA (GenScript) encoding a human IgD BCR with the variable region of the mAb 2134-1-A10 described in this study. To generate DQ2.5-sec-3-specific T cells, we sequenced the TCR of the human T-cell clone TCC1336.13^23^ using a template-switch anchored RT-PCR approach^35^. The TCR variable region in combination with mouse TCR α and β constant regions was then obtained as synthetic DNA (GenScript) that was subcloned into the pMIG II vector and used for transduction^36^.

### T cell-B cell collaboration assay

To assess the specificity of DQ2.5-glia-ω2-reactive (TCR737.30^16^) and DQ2.5-sec-3-reactive (TCR1336.13) BW58 T cells, 50,000 HLA-DQ2.5-expressing A20 B cells were incubated separately with the following peptides: deamidated 34mer harboring DQ2.5-glia-ω2 (QPQQPFPEQPQQPEQPF<u>PQPEQPFPW</u>QPEQPFPQ), deamidated 14mer harboring DQ2.5-Sec-3 (PQ<u>PFPEQPEQII</u>PQ), native ω-secalin 22mer (FPQQPQRPEQQFPQQPEQIISQ). B cells and peptides were incubated for 2 h at 37°C in 5% (v/v) FBS/RPMI, before 25,000 hybridoma T cells were added to the cultures. T-cell activation was assessed by ELISA detection of murine IL-2 in the supernatant after overnight incubation^34^. To assess peptide presentation by FPQQPEQII-reactive (2134-1-A10) and PLQPEQPFP-reactive (1002-1-E03^16^) A20 B cells, the cells were incubated with chymotrypsin-digested gluten that had been treated with recombinant TG2 to induce deamidation^23^ followed by addition of DQ2.5-glia-ω2-specific or DQ2.5-sec-3-specific hybridoma T cells as described above.

## ACKNOWLEDGEMENTS

We are thankful to Bjørg Simonsen and Marie K. Johannesen for technical assistance. The gluten protein database used for construction of the peptide array was compiled by Siri Dørum. Flow cytometry and cell-sorting experiments were conducted at the Flow Cytometry Core Facility, Oslo University Hospital. This work was funded by grants from Stiftelsen KG Jebsen (project SKGJ-MED-017) and the University of Oslo World-leading research program on human immunology (WL-IMMUNOLOGY).

## Supplemental Information

**Figure S1.**
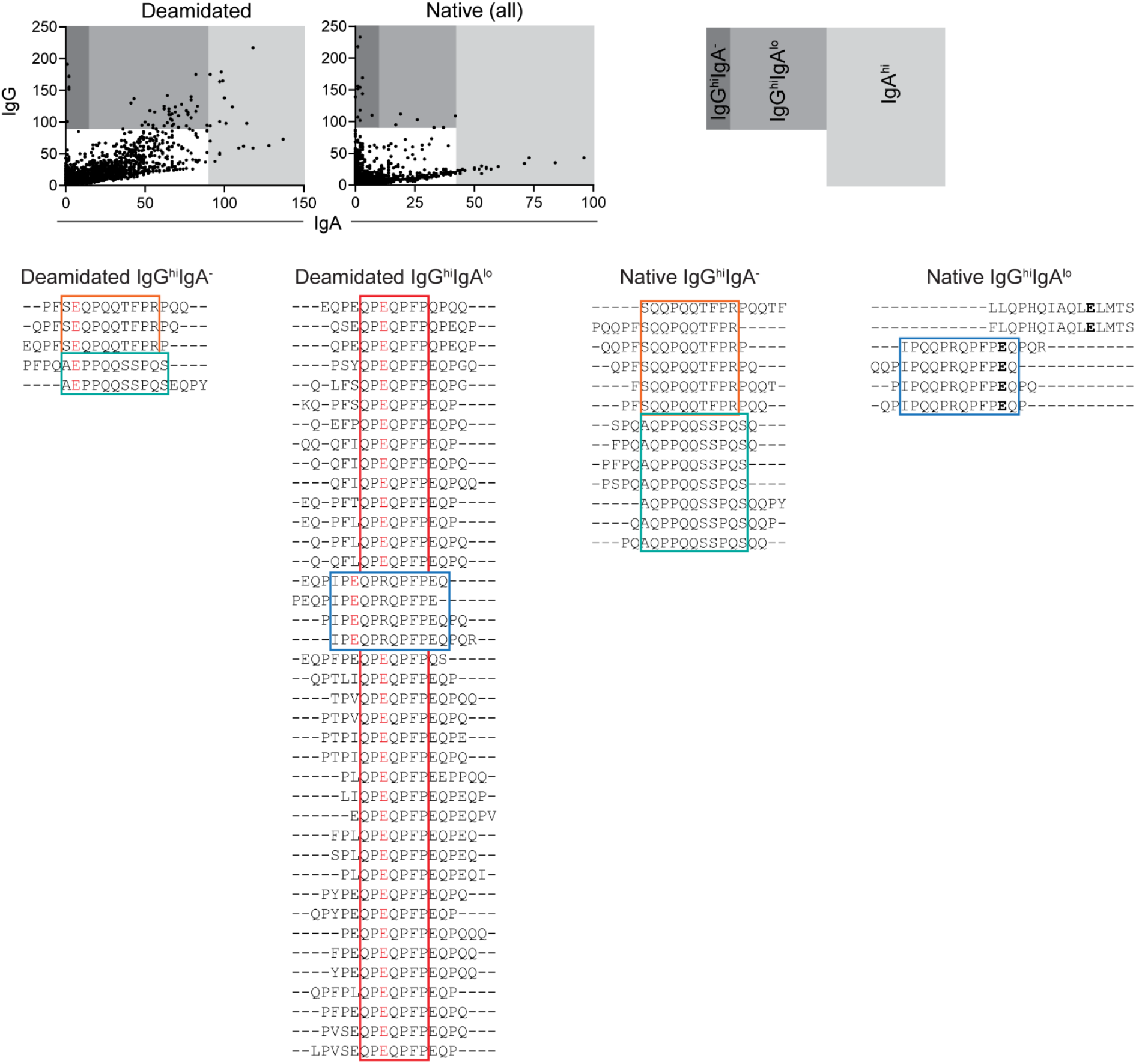
Gluten peptides with high serum IgG reactivity in CeD. Alignment of peptide sequences from deamidated and native gluten associated with high serum IgG reactivity in CeD patients. Peptides were either exclusively targeted by IgG or giving low IgA signals as indicated. Peptides with high IgA reactivity are shown in Fig. 1D. Colored boxes indicate recurring epitopes. Notably, the QPEQPFP epitope dominates both the IgA and the IgG response to deamidated gluten. All indicated epitopes contain PQQ or PEQ motifs. In addition, peptides associated with IgA reactivity all contain E, which is either introduced by deamidation (red) or present in native gluten (bold).

**Table S1.** Clinical data of CeD patients and non-celiac controls used for assessment of antibody reactivity in pooled sera and sorting of native-reactive gut plasma cells.

|  | Patient ID | HLA | Anti-TG2 IgA<sup>1</sup><br>(cut-off 5) | Anti-DGP IgG<sup>1</sup><br>(cut-off 20) | Marsh<br>score<sup>2</sup> | Diagnosis |
| --- | --- | --- | --- | --- | --- | --- |
| Antibody reactivity in pooled sera | CD1210 | DQ2.5 | 6.4 | 18 | 3B/1 | Untreated CeD |
|  | CD1212 | DQ2.5 | >100 | 83 | 3B-C | Untreated CeD |
|  | CD1214 | DQ2.5 | 27.7 | 13 | 3C/3B | Untreated CeD |
|  | CD1215 | DQ8 | >100 | 68 | 3B | Untreated CeD |
|  | CD1230 | DQ2.5 | 52.5 | 47 | 3B-C | Untreated CeD |
|  | CD1253 | DQ8 | >100 | >100 | 3B | Untreated CeD |
|  | CD1257 | DQ2.5 | >100 | 50 | 3B | Untreated CeD |
|  | CD1260 | DQ8 | 55.6 | 53 | 3B-C | Untreated CeD |
|  | CD1263 | DQ2.5 | >100 | >100 | 3C | Untreated CeD |
|  | CD1264 | DQ2.5 | 10.8 | 7 | 3C | Untreated CeD |
|  | CD1242 | —<sup>3</sup> | <1 | <5 | 0 | Non-celiac |
|  | CD1255 | —<sup>3</sup> | <1 | <5 | 0 | Non-celiac |
|  | Patient ID | HLA | Anti-TG2 IgA<sup>1</sup><br>(cut-off 7) | Anti-DGP IgG<sup>1</sup><br>(cut-off 7) | Marsh<br>score<sup>2</sup> | Diagnosis |
| Sorting of<br>plasma cells | CD2134 | DQ2.5 | >100<sup>4</sup> | 40<sup>4</sup> | 3C | Untreated CeD |
|  | CD5015 | DQ2.5 | 111 | 84 | 3B-C | Untreated CeD |
|  | CD5045 | DQ2.5/DQ8 | 15 | 18 | 0 | Potential CeD |
|  | CD5048 | DQ2.5 | >120 | 24 | 3B-C | Untreated CeD |

<sup>1</sup>Serum levels of TG2-specific IgA and deamidated gluten peptide (DGP)-specific IgG at the time of examination.
<sup>2</sup>Oberhuber G, Granditsch G, Vogelsang H. The histopathology of coeliac disease: time for a standardized report scheme for pathologists. *Eur J Gastroenterol Hepatol* 1999; **11**(10): 1185-1194.
Marsh score was determined in the third portion of the duodenum (d3). When two values are given, they represent histological findings in the duodenal bulb/d3.
<sup>3</sup>DQ2.5 and DQ8 negative.
<sup>4</sup>Cut-off 4 for anti-TG2 IgA and 20 for anti-DGP IgG

**Table S2. Serum antibody reactivity to native gluten peptides**

See Excel file.

**Table S3. Serum antibody reactivity to deamidated gluten peptides**

See Excel file.

**Table S4.**
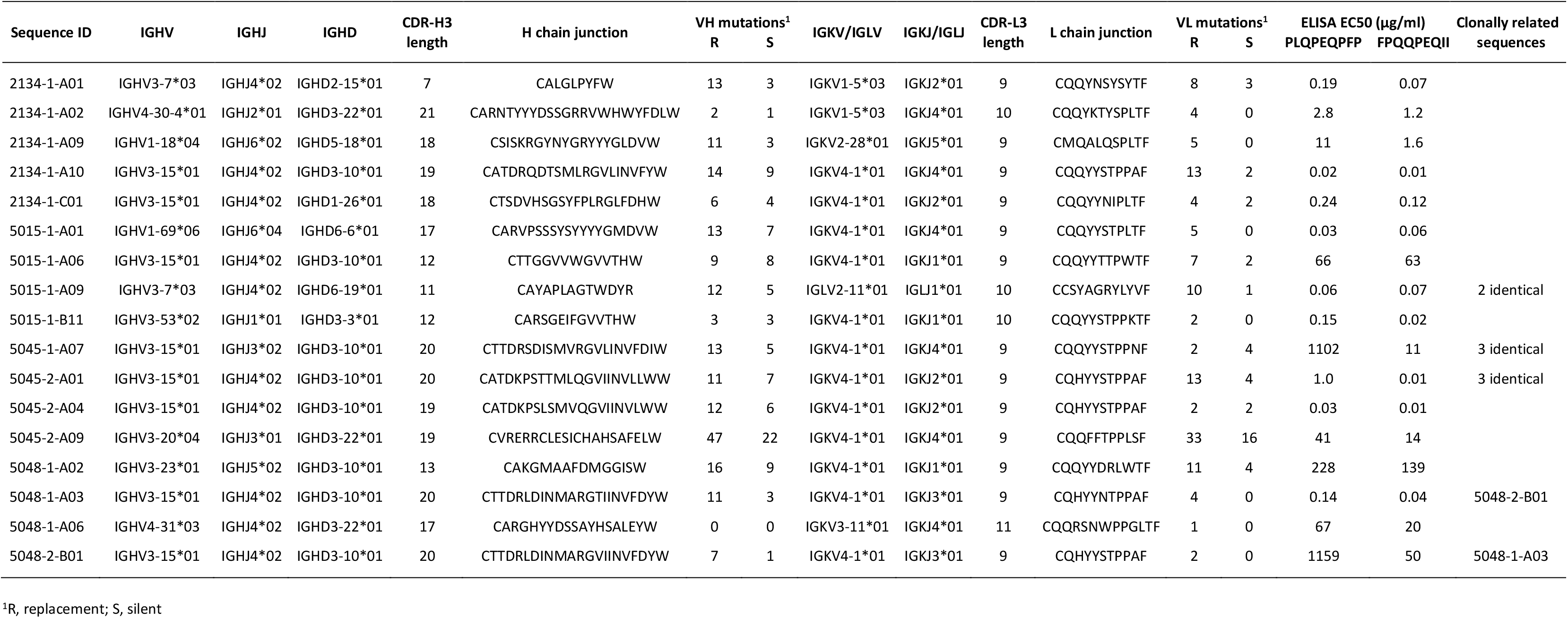
Sequence properties and reactivity of mAbs with confirmed binding to native gluten epitope.

